# Visual object categorization in infancy

**DOI:** 10.1101/2021.02.25.432436

**Authors:** Céline Spriet, Etienne Abassi, Jean-Rémy Hochmann, Liuba Papeo

## Abstract

Humans make sense of the world by organizing things into categories. When and how does this process begin? We investigated whether real-world object categories that spontaneously emerge in the first months of life match categorical representations of objects in the human visual cortex. Taking infants’ looking times as a measure of similarity, we defined a representational space where each object was defined in relation to others of the same or different categories. This space was compared with hypothesis-based and fMRI-based models of visual-object categorization in the adults’ visual cortex. Analyses across different age groups revealed an incremental process with two milestones. Between 4 and 10 months, visual exploration guided by saliency gives way to an organization according to the animate-inanimate distinction. Between 10 and 19 months, a category spurt leads towards a mature organization. We propose that these changes underlie the coupling between *seeing* and *thinking* in the developing mind.

## Introduction

Objects are the units of attention and perception; categories are the units of thought. We see objects (e.g., a given mushroom); but we think about objects primarily in terms of categories (e.g., *amanita muscaria*). By recognizing an object as member of a category, we understand what that object is and retrieve its visible (e.g., it is red with white spots) as well as its invisible properties (e.g., it is hallucinogenic). Categorization is, thus, the basis of inference and decision.

Objects can be categorized according to a virtually infinite number of perceptual and non-perceptual dimensions^1,2^. Insight on the most basic and general dimensions for object categorization in humans has been gained by studying how information is organized in the vast brain territory for visual object representation, which forms the occipitotemporal visual ventral stream.

Here, categories emerge from the topography of responses to visual objects, resolving into a large-scale organization that distinguishes between animate and inanimate objects, and crumbles in finer-grained distinctions between human *vs.* nonhuman animals, small *vs.* big (in terms of real-world size)^3,4^ natural *vs.* artificial objects^3,5–11^. Underneath this organization lays a mosaic of local *hot spots* of strong selectivity for stimuli such as faces, bodies and scenes^12–15^. Because of its organization and role in object recognition, the visual ventral stream is regarded as the interface between perception and cognition, forming the backbone for semantic categorization and representation of object and action knowledge in the rest of the brain^16^.

Besides the topography, categorical distinctions in the visual cortex also emerge from dissimilarities between distributed patterns of neural activity evoked by individual objects^7,17,18^. Thus, in visual areas, activity patterns recorded with functional MRI (fMRI) are more similar (i.e., less discriminable) for two animate objects (e.g., parrot and camel) than between an animate and an inanimate object (e.g., parrot and car). Visual object categories represented in the visual cortex prove behaviorally relevant, predicting the way in which individuals parse the visual world. For example, in visual search for a target-object among a set of distractors, people are faster to discriminate and find a target among objects of a different visual category (e.g., a cat among artificial objects), than among objects of the same visual category: search times increase as neural similarity between target and distractors increases^5^.

The organization of the human visual cortex by object categories appears to be a hallmark in the evolution of the primate brain: it is replicated in the visual cortex of monkeys^7,19^, and is resistant to variations of individual visual experience^20–23^. A similar organization across species and conspecifics with different environment and life-long visual experience, suggests a neural code optimized by evolution. This line of thinking encourages the hypothesis that object representation in visual cortex reflects biological constraints and dispositions^24^; as such, it would emerge early in life or even be present at birth.

There is initial evidence for signatures, or precursors, of neural specialization to object categories (faces, bodies, animals and scenes), in the visual cortex of newborns or young infants, using electroencephalography^25–28^ or fMRI^29,30^. Behavioral counterparts of those neural effects include early preference for faces or face-like stimuli over inverted faces^31–33^, for biological, over non-biological motion^34,35^, and for canonical, over distorted bodies^36–38^.

While preference implies discrimination between two objects, visual categorization entails the ability to use the canonical visual properties of a category (e.g., shape) to identify its members and keep them separate from other categories. By 4 months, infants are already able to do so: exposed to various exemplars of a category (e.g., cats), they exhibit a novelty effect, looking longer at an object of a new category than at a novel object of the same category^39,40^.

But, when do infants begin to see the visual world as adults do? Here, we investigated whether the categorical dimensions that drive the large-scale organization of the human visual cortex could account for the spontaneous emergence and development of real-word object categories in infancy. In particular, under the hypothesis that the structuring of visual object information toward an adult-like organization begins at birth^27,29,30^, we asked when such organization becomes functional so as to account for how infants explore the visual world.

We examined the development of visual object categorization in infancy, considering, in one experimental design, objects that have highlighted categorical representations in the visual cortex of human adults (and monkeys): animate *vs.* inanimate, human *vs.* nonhuman (animate), faces *vs.* bodies, natural *vs.* artificial inanimate, and real-world big *vs.* small (inanimate)^7^. Each of the above distinctions defines a categorization model, whereby a given (behavioral or physiological) correlate of object perception would be more similar for two objects of the same category than for two objects of different categories.

Using eye tracking, we recorded the most reliable and informative measure of infants’ cognition thus far: the looking behavior^41,42^. Infants of 4, 10 and 19 months viewed two objects at a time on a screen, while we measured the looking time towards either object. We took the looking time difference between two stimuli as a measure of discrimination, under the hypothesis that looking times for two objects seen for the first time would be more similar, the closer their visual representation is. This approach defined a model where each object category was represented in relation to the others –i.e., how similar/dissimilar it was from exemplars of the same and different categories. A model based on a relative measurement can be quantitatively compared with any model based on another relative measurement, whatever the source of the measurements (e.g., reaction times, neural activity)^43^. We compared the model of visual object representation emerging from the infants’ looking behavior, with synthetic (i.e., hypothesis-driven) and data-driven (i.e., fMRI-based) models reflecting visual object representation in the mature visual cortex. The current approach had previously allowed connecting data from brain-activity recording, behavioral measurement, and computational modeling. Here, we connected another branch, which is another step towards a unified theory of the origin and development of functional organization in the human brain.

## Results

### Experiment 1

Three groups of infants of 4 (*n* = 24), 10 (*n* = 24) and 19 months (*n* = 25) saw 36 pairs of images, each featuring an object from one of eight categories: human faces, human bodies, nonhuman faces, nonhuman bodies, natural-big and natural-small objects, artificial-big and artificial-small objects (hereafter, “big” and “small” refer to real-world size) (Fig. 1a). All subjects saw all possible combinations (Fig. 1b) of between-category and within-category pairs. For each infant, for each pair, we measured the absolute difference in looking times between the left and right images (differential looking time, DLT) *(Methods)*. DLTs were used to build a representational dissimilarity matrix (RDM), in which cells off the diagonal represented between-category comparisons and cells on the diagonal represented within-category comparisons (Fig. 2). Since different infants saw different exemplars for each category, group-averaged RDMs represented relationships–i.e., dissimilarities–between categories, rather than between individual objects.

**Figure 1.**
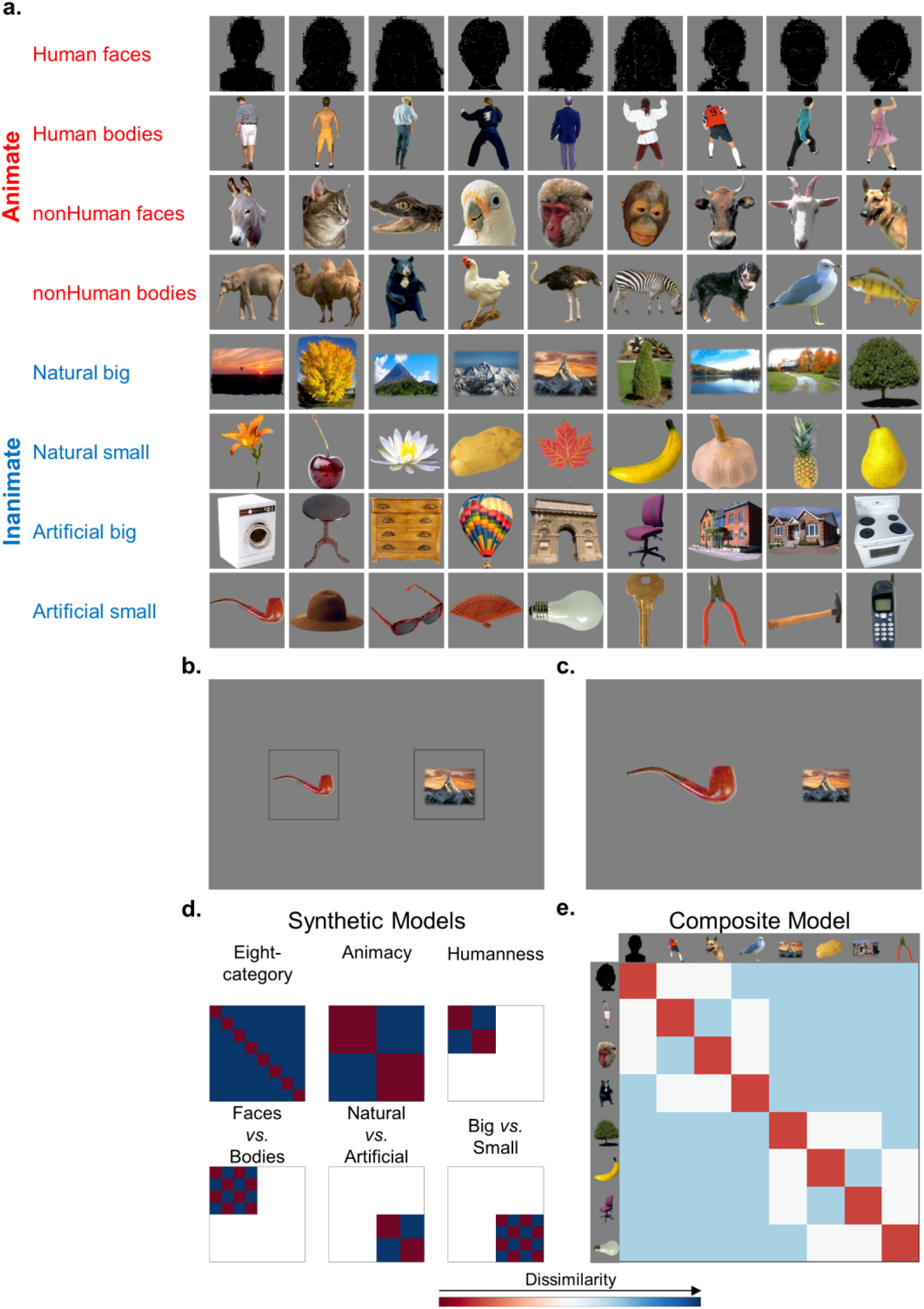
Stimuli, trials and hypothesis-based models of categorization considered in the design of Experiments 1-2. (**a)** Stimuli were 72 images depicting nine objects from each of eight different categories. Only silhouette of the face-stimuli used in the experiment are shown, because verification of consent of people in the pictures is incompatible with the rapid and automated nature of preprint posting. **(b)** In each trial of Experiment 1, two images were presented within two grey frames of identical size, on the right and on the left, equally distant from the center of the screen. (**c)** In each trial of Experiment 2, the image frame was removed and the image size was modified so that each object had the same number of pixels. (**d)** Hypothesis-driven (synthetic) models reflecting the categorical object representations tested in the current design. **(e)** The composite model reflecting the mean of the six synthetic models.

**Figure 2.**
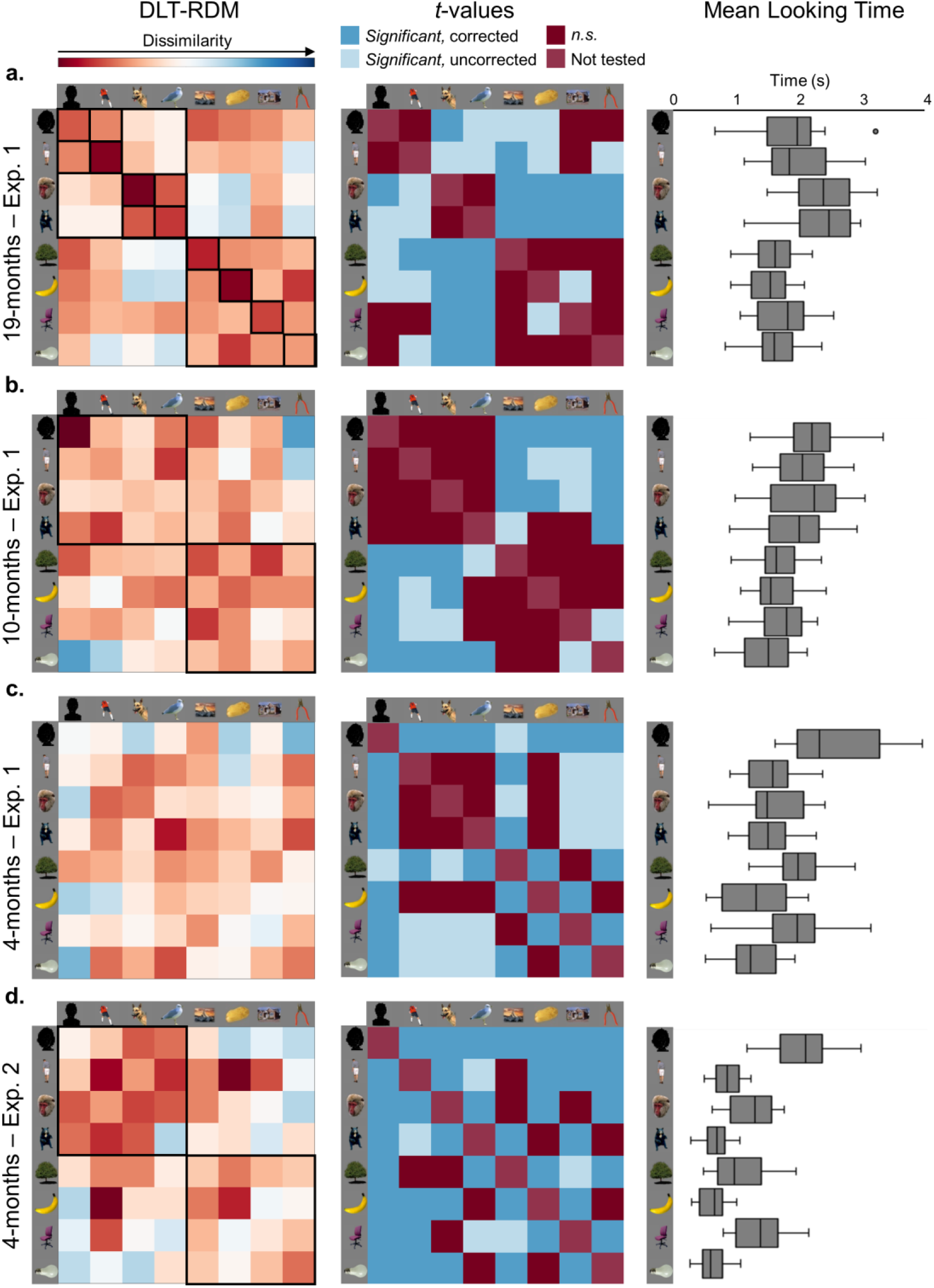
Results of representational similarity analysis and of the pairwise comparisons of mean looking times (MLT) between- and within-categories for each age group in Experiments 1-2. Left: Mean representational dissimilarity matrix (RDM) reflecting dissimilarities between- and within-categories in terms of differential looking times (DLTs). Black squares in the RDMs highlight categorization by animacy, humanness and by the eight categories in 19-month-olds **(a)**, categorization by animacy in 10-month-olds **(b)**, and in 4-month-olds of Experiment 2 **(d)**. **Center:** Matrix of *t*-values for each pairwise comparison between- or within-category with respect to the MLTs for 19- **(a)**, 10- **(b)** and 4-month-olds **(c)** in Experiment 1 and 4-month-olds in Experiment 2 **(d)**. Squares in dark blue denote significant effects; squares in lighter blue denote effects that did not survive the multiple comparison correction (trends); red squares denote non-significant (*n.s.*) or non-tested comparisons. **Right:** Distribution of MLTs in 19- (**a**), 10- (**b**) and 4-month-olds (**c**) of Experiment 1 and of 4 month-olds of Experiment 2 (**d**). Box-plots represent the minimum, the first quartile, the median, the third quartile and the maximum of the population distribution; outliers are denoted by dots (1 in the 19-month-old group).

#### Reference models of visual object categorization

Using representational similarity analysis^43^, we computed the relationship between RDMs based on infants’ DLT (DLT-RDMs) and models (i.e., RDMs) of visual object categorization in adults, defined with two independent approaches. The first approach defined a set of categorization models based on fMRI responses evoked in human adults, when viewing the same objects presented to infants. In the fMRI-based RDMs, pairwise between- and within-category dissimilarities reflected correlations between neural activity patterns *(Methods)*. Three RDMs were computed from activations in three broad regions of interest (ROIs; Fig. 3a-b) of the visual cortex (early visual cortex, EVC; ventral occipitotemporal cortex, VOTC; and lateral occipitotemporal cortex, LOTC), and at each location along the antero-posterior axis of the visual ventral stream (i.e., *vector-of-ROIs* analysis). The second approach defined six synthetic categorization models (RDMs) that may apply to the current stimulus-set: animate-inanimate (animacy model), human-nonhuman animates (humanness model), faces-bodies, natural-artificial inanimates, big-small inanimates, and eight-category model, where each category was defined as a category of its own, distinct from the other seven (Fig. 1d). In each cell of an RDM, the values 0 or 1 indicated dissimilarities within-category (lowest dissimilarity) and between-category (highest dissimilarity), respectively. As a model of visual object categorization in adults, a composite-RDM was obtained by averaging the above six models (Fig. 1e).

**Figure 3.**
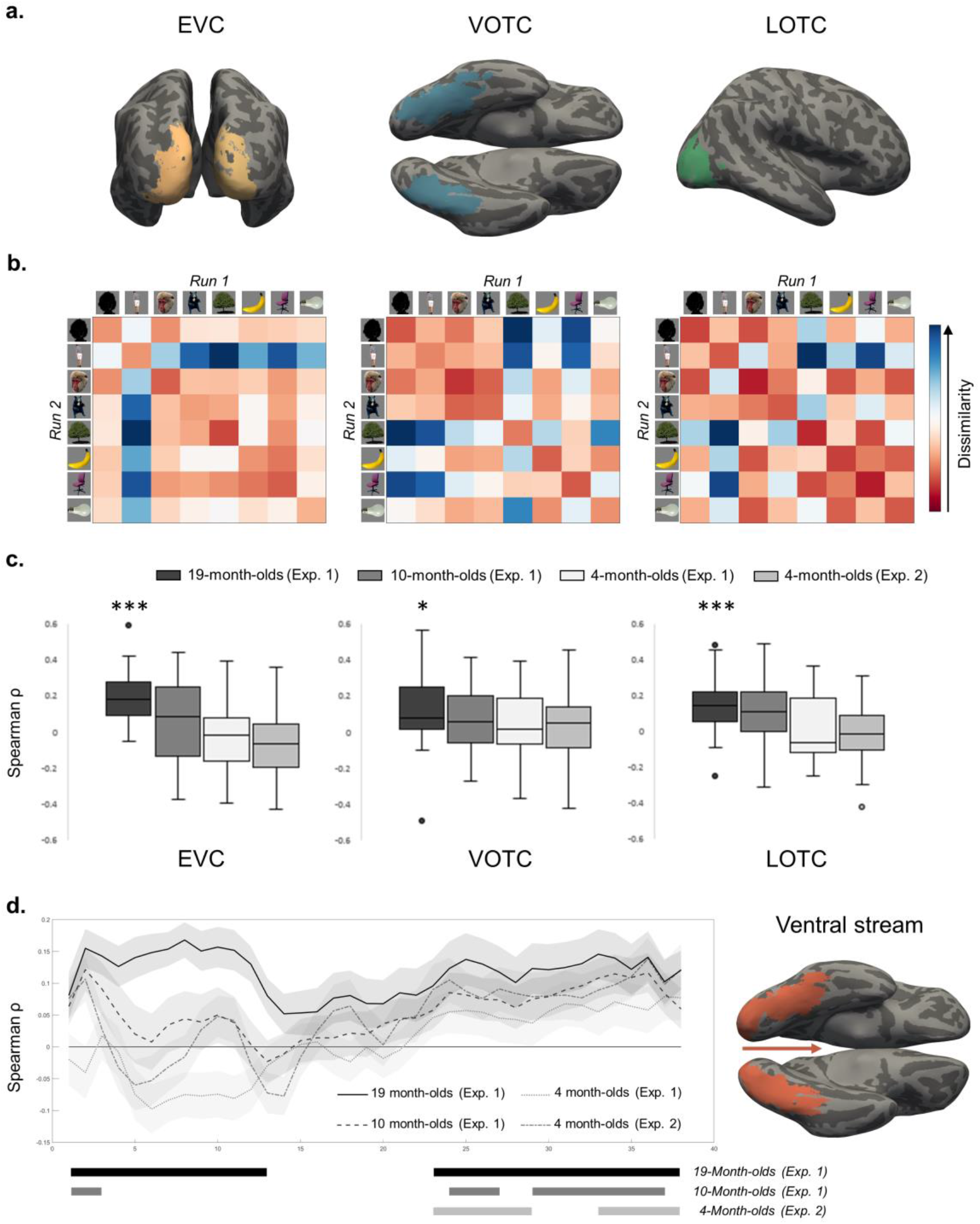
Relationship between infants’ looking behavior and visual object representation in the adults’ visual cortex. **(a)** Regions of interest (ROIs) in the adults’ brain: early visual cortex (EVC), ventral occipitotemporal cortex (VOTC) and lateral occipitotemporal cortex (LOTC). **(b)** Mean representational dissimilarity matrix (RDMs) reflecting relationships (i.e., dissimilarities) between object categories in terms of dissimilarities in the neural activity patterns evoked by viewing objects in the EVC, VOTC and LOTC of adults (fMRI-based RDMs). **(c)** Results of the representational similarity analysis between the mean fMRI-based RDM in each ROI and the RDMs based on differential looking times (DLT-RDM) of each infant in each age group of Experiments 1-2. Box-plots represent the minimum, the first quartile, the median, the third quartile and the maximum of the population distribution as well as outliers (dots); * *P* < 0.017; *** *P* < 0.0003. **(d)** Results of the representational similarity analysis between the infants’ DLT-RDM and the fMRI-based RDM derived each partition along the ventral visual stream. Solid bars represent clusters with significant correlation (above 0) for each age group of Experiments 1-2.

In addition to, or instead of, categorical information, infants’ look might be guided by physical properties of the stimuli such as size of the image on the retina^44,45^, elongation^46^ and color^46^. To assess systematic relations between looking times and visual features of the images irrespective of the category, we computed RDMs representing differences in size and elongation, relying on signed values, to appreciate the looking-time difference between two objects but also which one (the larger one; the more or less elongated one) was looked at the longer. A third RDM was computed to represent differences in the image color (Supplementary Fig. 1).

#### Nineteen-month-olds

The group-averaged DLT-RDM (Fig. 2a) showed an adult-like organization, as reflected by significant correlations with the composite-RDM, and the RDMs derived from the EVC, VOTC and LOTC (see statistics in Table 1; Fig. 3c). The vector-of-ROIs analysis showed that the DLT-RDM was maximally correlated with object-related responses in early visual areas (V1 to V3) and fusiform gyrus (*P*s < 0.001; Fig. 3d).

**Table 1.**
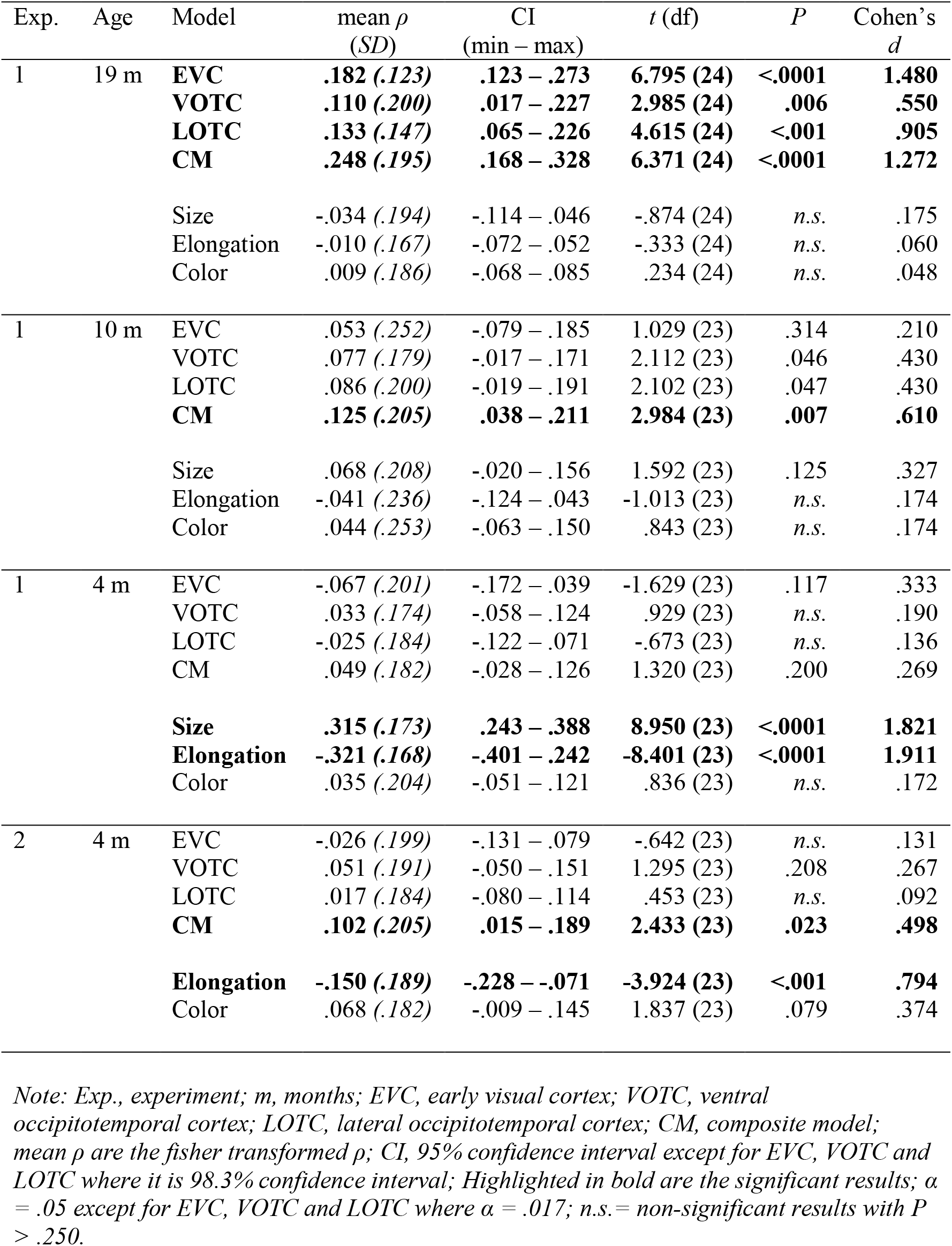
Results of representational similarity analysis reflecting the relationship of the infants’ DLT-RDMs with the fMRI-based RDMs, the RDM for the synthetic composite model of categorization and the RDMs based on size, elongation and color differences.

| Exp. | Age | Model | mean $\rho$<br>(*SD*) | CI<br>(min – max) | $t$ (df) | $P$ | Cohen's<br>$d$ |
| --- | --- | --- | --- | --- | --- | --- | --- |
| 1 | 19 m | **EVC** | **.182 (.123)** | **.123 – .273** | **6.795 (24)** | **<.0001** | **1.480** |
|  |  | **VOTC** | **.110 (.200)** | **.017 – .227** | **2.985 (24)** | **.006** | **.550** |
|  |  | **LOTc** | **.133 (.147)** | **.065 – .226** | **4.615 (24)** | **<.001** | **.905** |
|  |  | **CM** | **.248 (.195)** | **.168 – .328** | **6.371 (24)** | **<.0001** | **1.272** |
|  |  | Size | -.034 (.194) | -.114 – .046 | -.874 (24) | *n.s.* | .175 |
|  |  | Elongation | -.010 (.167) | -.072 – .052 | -.333 (24) | *n.s.* | .060 |
|  |  | Color | .009 (.186) | -.068 – .085 | .234 (24) | *n.s.* | .048 |
| 1 | 10 m | EVC | .053 (.252) | -.079 – .185 | 1.029 (23) | .314 | .210 |
|  |  | VOTC | .077 (.179) | -.017 – .171 | 2.112 (23) | .046 | .430 |
|  |  | LOTc | .086 (.200) | -.019 – .191 | 2.102 (23) | .047 | .430 |
|  |  | **CM** | **.125 (.205)** | **.038 – .211** | **2.984 (23)** | **.007** | **.610** |
|  |  | Size | .068 (.208) | -.020 – .156 | 1.592 (23) | .125 | .327 |
|  |  | Elongation | -.041 (.236) | -.124 – .043 | -1.013 (23) | *n.s.* | .174 |
|  |  | Color | .044 (.253) | -.063 – .150 | .843 (23) | *n.s.* | .174 |
| 1 | 4 m | EVC | -.067 (.201) | -.172 – .039 | -1.629 (23) | .117 | .333 |
|  |  | VOTC | .033 (.174) | -.058 – .124 | .929 (23) | *n.s.* | .190 |
|  |  | LOTc | -.025 (.184) | -.122 – .071 | -.673 (23) | *n.s.* | .136 |
|  |  | CM | .049 (.182) | -.028 – .126 | 1.320 (23) | .200 | .269 |
|  |  | **Size** | **.315 (.173)** | **.243 – .388** | **8.950 (23)** | **<.0001** | **1.821** |
|  |  | **Elongation** | **-.321 (.168)** | **-.401 – .242** | **-8.401 (23)** | **<.0001** | **1.911** |
|  |  | Color | .035 (.204) | -.051 – .121 | .836 (23) | *n.s.* | .172 |
| 2 | 4 m | EVC | -.026 (.199) | -.131 – .079 | -.642 (23) | *n.s.* | .131 |
|  |  | VOTC | .051 (.191) | -.050 – .151 | 1.295 (23) | .208 | .267 |
|  |  | LOTc | .017 (.184) | -.080 – .114 | .453 (23) | *n.s.* | .092 |
|  |  | **CM** | **.102 (.205)** | **.015 – .189** | **2.433 (23)** | **.023** | **.498** |
|  |  | **Elongation** | **-.150 (.189)** | **-.228 – -.071** | **-3.924 (23)** | **<.001** | **.794** |
|  |  | Color | .068 (.182) | -.009 – .145 | 1.837 (23) | .079 | .374 |

*Note: Exp., experiment; m, months; EVC, early visual cortex; VOTC, ventral occipitotemporal cortex; LOTc, lateral occipitotemporal cortex; CM, composite model; mean $\rho$ are the fisher transformed $\rho$ ; CI, 95% confidence interval except for EVC, VOTC and LOTc where it is 98.3% confidence interval; Highlighted in bold are the significant results; $\alpha = .05$ except for EVC, VOTC and LOTc where $\alpha = .017$ ; *n.s.*= non-significant results with $P > .250$ .*

Next, we asked which of the six categorical models underlying the composite-RDM best represented the infants’ DLT-RDMs. A stepwise linear regression (α_corrected_: 0.0083, two-tailed) showed an effect of the eight-category model (mean *β* = 0.090; 99.17% CI [100*(1-0.0083)] = 0.026 – 0.155; *t*(24) = 4.023, *P* < 0.001; *d* = 0.804), animacy model (mean *β* = 0.077; 99.17% CI = 0.014 – 0.139; *t*(24) = 3.514, *P* = 0.002; *d* = 0.702), and humanness model (mean *β* = 0.133; 99.17% CI = 0.009 – 0.256; *t*(24) = 3.091, *P* = 0.005; *d* = 0.618) (for all other regressors: *P*s *>* 0.07; Supplementary Table 1).

In a new analysis, we considered another measure of categorization: for the six categorization models, we tested whether average between-category DLTs were higher than average within-category DLTs (α_corrected_: 0.0083, one-tailed). Confirming previous results, we found that this was the case for the eight-category model (*M_difference_* = −0.135; 99.17% CI = -inf – −0.091; *t*(24) = −7.967, *P* < 0.0001; *d* = 1.588), animacy model (*M_diference_* = −0.095; 99.17% CI = -inf – −0.041; *t*(24) = −4.481, *P* < 0.0001; *d* = 0.896) and humanness model (*M_difference_* = −0.158; 99.17% CI = -inf – −0.060; *t*(24) = −4.16, *P* < 0.001; *d* = 0.832), but not for the other models (Supplementary Table 2).

Following evidence of categorization based on the eight-category model, we asked which of the eight categories the infants could indeed represent. Separately for each of the eight categories, we tested whether average within-category DLTs were lower than average between-category DLTs (*t*-tests; α_corrected_: 0.0063, one-tailed), and found that, in addition to animates, inanimates, humans and nonhumans, 19-month-olds showed to represent the subordinate categories of human bodies (*M_difference_* = −0.195; 99.37% CI = -inf – −0.060; *t*(24) = −3.902, *P* < 0.001, *d* = 0.78), nonhuman bodies (*M_difference_* = −0.160; 99.37% CI = -inf – −.014; *t*(24) = −2.956, *P* = 0.004, *d* = 0.603), nonhuman faces (*M_difference_* = −0.216; 99.37% CI = -inf – −.094; *t*(24) = −4.763, *P* < 0.0001, *d* = 0.953) and natural-small objects (*M_difference_* = −0.179; 99.37% CI = -inf – −0.076; *t*(24) = −4.727, *P* < 0.0001, *d* = 0.986) (all other *P*s > 0.024).

Difference between images in color, size or elongation did not account for the infants’ looking behavior (Table 1), suggesting priority of categorical information, over more general physical differences, in the 19-month-olds’ processing of visual objects.

Finally, we assessed possible preferences, considering the mean looking times (MLTs) toward each category, averaged across trials and subjects. A one-way repeated-measures ANOVA showed an effect of Category (*F*(7,168) = 16.259, *P* < 0.0001; η^2^ =0.790), which reflected a preference (i.e., longer looking times) for animate (*M* = 2.103 s ± 0.344) over inanimate categories (*M* = 1.598 s ± 0.247; *M_difference_* = 0.505; 95% CI = 0.367 – 0.643; *t*(24) = 7.551, *P* < 0.0001; *d* = 1.509), and for nonhuman animals (*M* = 2.317 s ± 0.463) over humans (*M* = 1.889 s ± 0.452; (*M_difference_* = −0.428; 95% CI = −0.676 – −0.179; *t*(24) = −3.552, *P* = 0.002; *d* = 0.711) (see Supplementary Results 3 for details). The matrix representing *t* values for each pairwise comparison of MLTs remarkably replicated the structure of the DLT-RDM (Fig. 2a; Supplementary Table 3 for *t*- and *P*-values), showing categorization and discrimination based on animacy and humanness.

#### Ten-month-olds

The looking behavior of 10-month-olds (Fig. 2b) was significantly correlated with the composite-RDM of adult categorization as well as with fMRI-based RDMs reflecting object-related responses in selective aspects of the adults’ visual ventral stream. Further analyses showed that objects were principally categorized by animacy.

More precisely, although correlations of the infants’ DLT-RDM with activations in the broad ROIs (EVC, VOTC and LOTC) did not reach the significance level (Table 1; Fig. 3c), the vector-of-ROIs analysis showed correlation with RDMs derived from the early visual cortex (V1) and fusiform gyrus (*P*s < 0.001; Fig. 3d). The DLT-RDMs also correlated with the synthetic composite-RDM. Infant’s behavior was not explained by differences in visual feature such as color, size or elongation (Table 1).

Which of the six models underlying the composite-RDM best represented the infants’ behavior? A stepwise linear regression showed correlation of the infants’ DLT-RDM with the animacy model only (α_corrected_: 0.0083, two-tailed; mean *β* = 0.059; 99.17% CI = 0.002 – 0.117; *t*(23) = 2.981, *P* = 0.007; *d* = 0.608; for all other regressors: *P*s *>* .036; Supplementary Table 1). Consistent with this finding, average within-category DLTs were significantly lower than average between-category DLTs for the animacy model (*M_difference_* = −0.061; 99.17% CI = -inf – −0.007; *t*(23) = −2.919, *P* = 0.004; *d* = 0.592; α_corrected_: 0.0083, one-tailed), but not for the other models (Supplementary Table 2).

The analysis of the MLTs revealed an effect of Category (one-way repeated-measures ANOVA: *F*(7,161) = 9.422, *P* < 0.0001; η^2^ =0.808), reflecting a preference (i.e., longer looking times) for animate (*M* = 2.044 s ± 0.316) over inanimate categories (*M* = 1.601 s ± 0.263; *M_difference_* = 0.443; 95% CI = 0.310 – 0.576; *t*(23) = 6.901, *P* < 0.0001; *d* = 1.409; see Supplementary Results 3). Relationships between categories computed on the MLTs replicated the structure of the DLT-RDM, showing that 10-month-olds categorized objects based on animacy, and exhibited a preference for animate objects (Fig. 2b; Supplementary Table 3).

#### Four-month-olds

Unlike older infants, 4-month-olds showed no evidence of categorization; they looked longer at human faces and big-inanimate objects or, otherwise, at the larger and less elongated of two images on the screen.

More precisely, the infants’ DLT-RDM did not match the organization of object-related information in any broadly defined ROIs of the adults’ visual cortex (Table 1; Fig. 3c), or in any smaller partition of the visual ventral stream (vector-of-ROIs analysis; Fig. 3d). No correlation was found with the composite model of adult categorization (Table 1) or with any of the six underlying models (stepwise linear regression: all *t*s < 1, *n.s.;* Supplementary Table 1). For none of the categorization models were DLTs larger for between-category, than within-category comparisons (*P*s > 0.065; Supplementary Table 2). Instead, infants’ DLT-RDM correlated positively with the RDM based on image size, and negatively with the RDM based on elongation (Table 1). No correlation was found with the color model.

A one-way repeated-measures ANOVA on the MLTs showed an effect of Category (*F*(7,161) = 22.970; *P* < 0.0001; η2 =0.802), which was driven by a preference for human faces over all other categories (α_corrected_: 0.0018; all *P*s < 0.001; Fig. 2c; Supplementary Table 3). Within the inanimate categories, infants looked longer at big over small objects, whether artificial or natural (*Ps* < 0.0001). We note that the two most preferred categories (human faces and big inanimates) were those with the largest image size (> 15000 pixels) and the least elongated shape (see Supplementary Results 4; Supplementary Fig. 2). Thus, size and/or elongation, rather than object identity, could explain object preferences in 4-month-olds. In line with this, the MLTs computed for each image across subjects, correlated positively with image size (*ρ* = 0.515, *P* < 0.0001), and negatively with elongation (*ρ* = −0.531, *P* < 0.0001). That is, the larger the image or the less elongated the shape, the longer the looking time. Given this result, with a new stepwise linear regression, we reassessed the relationship of the DLT-RDM with the six categorical models, after removing the variance explained by size and elongation. Yet, no model accounted for the remaining variance (all *Ps* > 0.263; see Supplementary Results 4; Supplementary Table 4). Congruently, we found no evidence of categorization comparing average within- and between-category DLTs (all *Ps* > 0.175; see Supplementary results 4; Supplementary Table 5).

#### Comparison between groups

The above analyses showed that categorization by animacy emerged by 10 months, while categorization by humanness, and additional categories of the eight-category model emerged by 19 months. Additional between-subjects analyses confirmed the differences between age groups, with respect to object categorizations by animacy, humanness and by the eight-category model (see Supplementary Results 5).

### Experiment 2

In Experiment 1, 4-month-olds showed no evidence of categorization, but a preference for human faces and big objects, which might be explained by physical properties such as image size and elongation –i.e., a tendency to look at the larger/less elongated image on the screen. Here, we asked whether a prominent preference for certain physical properties might have overshadowed categorical effects. We tested a new group of 4-month-olds (*n* = 24) with the same images of Experiment 1, but all matched for size (i.e., number of pixels; Fig. 1c). The size, but not the elongation, was modified because only the former can change without affecting object identity or recognizability. Results confirmed the preference for human faces and big-inanimate object, but also showed that, when size was no longer available to discriminate between two stimuli, categorization by animacy emerged in 4-month-olds too.

Specifically, the infants’ DLT-RDMs (Fig. 2d) correlated with the composite-RDM, and the fMRI-based RDMs extracted from the anterior fusiform gyrus (*P* < 0.001; Fig. 3d) in the vector-of-ROIs analysis (correlations with broad ROIs did not reach significance; see Table 1). There remained a significant negative correlation with the elongation model, and no correlation with the color model (Table 1).

Of the six synthetic models that contributed to the composite-RDMs, infants’ behavior was best represented by the animacy model (stepwise linear regression, α_corrected_: 0.0083, two-tailed; mean *β* = 0.074; 99.17% CI = 0.016 – 0.132; *t*(23) = 3.70; *P* = .001; *d* = 0.755; for all other regressors, *Ps >* 0.12; Supplementary Table 1). The comparison between within-category and between-category DLTs confirmed the above results, showing lower within-category than between-category DLTs for the animacy model (α_corrected_: 0.0083, one-tailed; *M_difference_* = −0.076; 99.17% CI = -inf – −0.021; *t*(23) = −3.583, *P* < 0.001; *d* = 0.731), but not for the other models (Supplementary Table 2).

A one-way repeated-measures ANOVA on the MLTs showed a significant effect of Category (*F*(7,161) = 59.466; *p* < 0.0001; η^2^ = 0.869), which reflected a preference for human faces over all other categories (α_corrected_: 0.0018; all *Ps* < 0.0001; see Supplementary Results 3; Supplementary Fig. 3) and for big- over small-inanimate objects (all *Ps* < .001). Thus, the preference for human faces and big objects, which were the largest objects in Experiment 1, remained despite matching images for size. Moreover, the average MLTs for individual images were negatively correlated with elongation (*ρ* = −0.396, *P* < 0.001), confirming the elongation bias: infants looked longer at less elongated shapes. Given the last result, we reassessed the correlation of the DLT-RDM with the six categorical models, after removing the variance explained by elongation. Again, the animacy model was the only significant regressor (α_corrected_: 0.0083; mean *β* = 0.064; 99.17% CI = 0.008 – 0.121; *t*(23) = 3.282; *P* = 0.003; *d* = 0.667; for all other regressors, *Ps >* 0.126; Supplementary table 4). Categorization by animacy was confirmed by higher between- than within-category DLTs for the animacy model only (α_corrected_: 0.0083, one-tailed; *M_difference_* = −0.068; 99.17% CI = -inf – −0.015; *t*(23) = −3.314; *P* = 0.002; *d* = 0.673; for all other comparisons *P* > 0.111; see Supplementary results 4; Supplementary Table 5).

## Discussion

Categorization is the mechanism through which the human mind makes sense of the environment by organizing the *things* of the world in categories. Categorization begins at a young age, with the ability to appreciate perceptual similarities between objects (or facts), and acquires refinement with knowledge and language acquisition. What type of real-world object categories infants can represent before developing a sizable lexicon and a rich system of knowledge about the world? We considered the hypothesis that the early stages of spontaneous visual object categorization are guided by the dimensions that structure object representations in the visual cortex of the primate brain.

Our findings demonstrate that early visual object categorization along the fundamental dimensions represented in the human visual cortex, is an incremental process with two milestones. The first, between 4 and 10 months, establishes the transition from an exploration of the environment guided by general visual saliency to an organization based on the broad animate-inanimate categorical distinction; the second, between 10 and 19 months, presents a spurt of visual object categories towards mature organization.

All the categorization effects that we observed analyzing looking time differences between objects were paired with preference effects, as indexed by mean looking times. Thus, when infants showed to categorize objects by animacy, they also looked longer at animate than inanimate objects; or, when they showed to categorize objects by humanness, they also looked longer at nonhuman than human animals. Note that, with our approach, we could observe categorization (e.g., animate *vs.* inanimate objects) as long as within-category differences were lower than between-categories differences, and even if some infants preferred one category, and others, the other category, thus yielding null group preference effect. The co-occurrence of preference and categorization demonstrates that the earliest visual categories to emerge in infancy are those that are important enough to give rise to a hierarchy of preferences.

### Within 4 months

Four-month-olds showed no evidence of categorization based on any of the categorical dimensions considered here, when image size allowed discriminating between two images on the screen (Experiment 1). Accordingly, their looking behavior did not match visual object representation in any sector of the adults’ visual cortex. Infants’ look, however, was not random. They looked longer at the larger image and the less elongated object on the screen, with size and elongation differences predicting looking time differences. Moreover, mean looking times revealed preference for human faces and real-world-size big (*vs.* small) inanimate objects.

The preference for human faces, extensively documented in 4-month-olds and younger infants^31–33^, has been explained by the detection of the characteristic eyes-mouth configuration, and/or the characteristic iris-pupil-sclera contrast of human eyes^31^. However, if performance here reflected detection of those features, infants would have shown categorization of human faces, as all our human faces carried those features. Instead, preference for faces occurred without categorization (i.e., within-category DLTs were as high as between-categories DLTs). This suggests that, in processing two faces, infants focused on individual- rather than category-level features, which allow for individuation, rather than categorization^47^.

As with human faces, the preference for real-world big (*vs.* small) objects emerged without evidence of categorization (i.e., within-category and between-categories DLTs were comparable). The effect of real-world object size at such a young age is unprecedented and open to multiple interpretations. In effects, scholars are debating a proper characterization of the big-small object distinction, which might relate to differences in perceptual properties (e.g., texture, spatial frequency)^8^ and/or to behavior-relevant properties of the objects^48^. The current results add a new piece to the puzzle, showing an early asymmetry between the two object categories.

In the adult brain, big objects, which typically function as landmarks (e.g., buildings and trees), are represented in the ventral aspect of the visual cortex, adjacent to place- and scene-specific areas. Small objects, which are graspable objects, are represented in more lateral aspects of the occipitotemporal cortex, also hosting areas for the representation of tools and actions^4^. Areas of the scene- and place-specific network are functionally interconnected already in the first weeks of life^30^, and responds strongly to scenes in 4-to-6-month-olds^29^. In contrast, at 4 months, infants are unable to grasp objects, showing immaturity of the networks that control hand movements and hand-object interaction. Interest in graspable objects increases during the first year of life^45,46^, as infants develop grasping skills^49^. Consistent with this trajectory, we found that by 10 months, the preference for big over small objects had disappeared. Thus, different developmental trajectories of different networks in the visual cortex might contribute to object distinctions captured by the effect of real-world size.

### From 4 to 10 months

When size was no longer available to discriminate between two images on the screen (Experiment 2), 4-month-olds continued to show preference for human faces and big (*vs.* small) objects, but they also showed categorization by animacy. Thus, the animate-inanimate categorization was functional at 4 months, but was overshadowed by the physical features, such as size, making an object more visible, independently from the category. By 10 months, infants showed to overcome the importance of low-level visual features in favor of categorical information: categorization by animacy emerged despite differences in image size. Moreover, by 10 months, the preference for human faces and big (*vs.* small) objects had given way to interest in the broader category of animate entities.

Thus, the looking behavior of both 10-month-olds in Experiment 1, and 4-month-olds in Experiment 2 revealed categorization by animacy and matched the cortical organization of object-related information recorded from anterior (temporal) aspects of the visual ventral stream in adults. Yet, the shift from prioritizing image size to prioritizing categorical information suggests a change between 4 and 10 months, which might result from increased integration of information on low-level features in earlier visual areas with categorical information in higher-level visual areas. Promoter of this developmental change could be the myelination of fiber tracts connecting distant areas^50^, which begins around 4 months in the occipital lobe and continues later, through the temporal lobe^51^.

Animacy is the earliest categorical distinction of visual objects in infancy. This means that representation of animate entities is not an extension of the representation of conspecifics (e.g., see the “human-first hypothesis”^47,52^). Infants would rather start with a broad, underspecified representation of what animates look like, which might function as a coarse “life detector” to identify conspecifics as well as predators and preys^53^. The animate-inanimate distinction thus lays the foundation for subtler categorical distinctions, while possibly setting conditions for domain-specific processes of naïve psychology^54,55^ *vs.* naïve physics^56,57^.

### From 10 to 19 months

We observed a second developmental change between 10 and 19 months, when infants showed to represent the categories of animate and inanimate objects, but also of humans, nonhuman animals, human bodies, nonhuman bodies and nonhuman faces. The spurt of categories by 19 months represents another step towards the model of mature visual object representation addressed here.

While in 4- and 10-month-olds, categorization limited to two categories was associated with object-related responses in the most anterior aspects of the visual cortex only, 19-month-olds’ behavior correlated with object-related responses across the broader visual system of adults (from early visual cortex to ventral and lateral higher-level areas and from posterior to anterior regions along the ventral stream). This fact suggests that the ability to form new visual categories, from superordinate (e.g., animate-inanimate) to basic and subordinate (e.g., human *vs.* nonhuman bodies), involves the progressive recruitment of more and more feature spaces distributed over the visual cortex, and representing features with different complexity (e.g., from low-to mid-level features). Again, myelination of tracts connecting distant regions might be pivotal in this process^50,51^. In the second year of life, categorization might also be supported by language development. Verbal labeling and communication of information about objects promote and shape the formation of new categories and, in some models, govern the transition from perceptual to conceptual categories^52,58–62^. It however remains possible that early visual object categorization is a perceptual matter, emerging as a consequence of functional properties of neurons in visual areas, brain maturation and early visual experience.

### Conclusions

We have shown that infants initiate their exploration of the visual world giving priority to images that are more visible –i.e., the larger ones. By 10 months, they show to learn that categorical information is more important than general physical properties. The first act of visual object categorization divides the world into animate and inanimate entities. Other categories represented in the visual cortex emerge later, by 19 months, perhaps because of a greater demand for visual experience and/or different maturation latencies in different visual areas. As visual categories multiply, infants’ behavior correlates with neural activity in ever-larger aspects of the adult visual cortex. Integration through growing connections within category-specific networks and between distant visual areas could be the driving force of this process. Increasing representation and reliance on visual categorical information in the first years of life may signal the coupling between *seeing* and *thinking*.

## Material and methods

### Eye-tracking study

#### Participants

The study involved 97 infants in total. Experiment 1 involved 24 infants of 4 months (11 females; mean age: 4 months, 15 days; range: 4:0-4:24), 24 infants of 10 months (8 females; mean age: 10:26; range: 10:1-11:30) and 25 infants of 19 months (11 females; mean age: 19:5; range: 18:1-20:1). Experiment 2 involved 24 infants of 4 months (14 females; mean age: 4:17; range: 4:3-5:0). The sample size of 24 was arbitrarily chosen for the initial group of 19-month-olds. Next, we verified that it was superior to the minimal sample size (*n* = 18) required to obtain the smallest categorical effect found in 19-month-olds (human *vs.* nonhuman: *d*_Cohen_= 0.6182, power = .80, α = .05; GPower 3.1), and kept it constant across groups. We continued testing until we reached 24 participants per group. The last 19-month-old infant had a twin; parents asked to test him too and we kept him in the sample. Fifty additional infants were tested in Experiments 1-2, but discarded *(Analysis)*. Infants were tested in the Babylab of the Institute of Cognitive Sciences Marc Jeannerod (Bron). Parents received travel reimbursement and gave informed consent before participation. The study was approved by the local ethics committee (CPP sud-est II).

#### Stimuli

We selected 72 total color photographs of isolated real-world objects, from publicly available sets^63^ or from the web. For Experiment 1, objects were superimposed on a grey background and scaled to fit a 350×350 pixels black frame. The final set of images consisted of nine exemplars for each of eight categories (human and nonhuman faces and bodies; natural and artificial big and small objects). Human faces were all female faces. For Experiment 2, all objects were resized to have the same number of pixels (54135) without grey background (Fig. 1c for an example).

#### Procedure

Infants sat on their parent’s lap, ~60 cm away from a Tobii Eye-tracker T60XL screen, in a dark room. Parents were instructed to keep their eyes close throughout the experiment. The experiment began after the calibration for eye-tracking, and consisted of 36 trials. In a trial, two images were presented for 5 s on the left and right side of the screen, equally distant from central fixation (Fig. 1b and c for examples). Each infant saw a unique set of pairs including all 28 possible between-category combinations and eight within-category combinations. The experiment ended after 36 trials (~3 minutes), or because the infant expressed discomfort, or stopped looking at the screen. Stimulus presentation and data recording were controlled through PsyScopeX (http://psy.cns.sissa.it).

#### Analyses

On the eye-tracking screen, we defined two areas-of-interest overlapping with the locations of the two images. Areas-of-interest were two 350×350 pixels squares in Experiment 1, and two masks encompassing all non-background pixels in Experiment 2. For every trial, we computed the cumulative looking times in each area-of-interest.

Only trials with ≥ 1 s of look within the areas-of-interest were considered valid. For the analyses, we discarded infants with less than 27 valid trials (3/4 of total trials) or with a strong side bias (i.e., fixation on the same side for > 80% of the experiment duration). In the final analysis of Experiment 1, 4-month-olds contributed, on average, 35 ± 1 trials, 10-month-olds, 34 ± 2 trials, and 19-month-olds, 34 ± 2 trials. In Experiment 2, 4-month-olds included in the final analysis contributed on average 29 ± 2 trials. Of all the infants tested in Experiment 1, exclusion criteria led to discard 24 because of insufficient data (13 4-month-olds, 7 10-month-olds and 4 19-month-old) and one 4-month-old because of a side bias. Twenty-five tested in Experiment 2 were discarded because of a side bias (*n* = 2) or insufficient data (*n* = 23). However, stimulus presentation differed between Experiments 1 and 2. In Experiment 1 areas-of-interest were fix, squared areas delimited (and highlighted) by a black frame, which contained object and background. In Experiment 2, areas-of-interest overlapped with the object’s contours without background and frame. In sum, in Experiment 2, stimuli might have been less visually salient and, therefore, the exclusion criteria, more stringent, than in Experiment 1, yielding higher attrition rate. To address this, we carried out additional analyses adopting more lenient criteria in Experiment 2, to reach an attrition rate closer to that of Experiment 1. This analysis confirmed all our results (Supplementary results 6).

For each infant, for each trial, one DLT was computed as the difference in the cumulative looking time (LT) between the right and the left area-of-interest divided by the sum of the two (i.e., the total time the infant attended to the areas of interest): (LT_right_-LT_left_)/(LT_right_+LT_left_). Absolute and signed DLTs values were entered in absolute and signed DLT-RDMs, respectively. Values on the diagonal (within-category DLTs) and off-diagonal values in one half of the DLT-RDM (between-category DLTs) were used for analysis.

#### Category effects

Separately for each experiment, for each group, we performed RSA to correlate the absolute DLT-RDMs with each of six categorization models (animacy, humanness, face-body, natural-artificial, big-small, and eight-category model) and the composite model of adult categorization, reflecting the average of the above six models. Each model defined an RDM, where cells had value of 0, 1 or 0.5 corresponding to within-category comparisons (i.e., lowest dissimilarity), between-category comparisons (i.e., highest dissimilarity), and comparisons irrelevant for a given categorization, respectively. For example, the humanness model had 0 for human-human (e.g., human face-human body) and nonhuman-nonhuman comparisons (e.g., cow face-elephant body), 1 for human-nonhuman comparisons (e.g., human body-camel body), and 0.5 for irrelevant comparisons (e.g., artificial-small object-natural large object).

First, we computed the correlation between the composite-RDM and the DLT-RDM of each infant. Individual *Spearman* correlation coefficients *ρ* for a group of infants were fisher-transformed and tested against chance-level 0 (*t*-test). Then, we performed a stepwise linear regression analysis for each infant, with the above six categorical models as regressors. For each regressor, the distribution of coefficients *β* in a group was compared against chance (*t*-test). Categorization was further addressed by assessing whether, for each model, average within-category DLTs were higher than average between-category DLTs (*t*-tests, one-tailed). All above analyses were computed considering the DLTs over the total 5 s trial duration.

#### Effects of general properties of the images

In Experiment 1, for each image, we computed a score for: 1) size (i.e., total number of pixels [350×350] minus number of background pixels); 2) shape-elongation (i.e., height-to-width ratio with ratio tending to 1 indicating lowest elongation), and 3) color (i.e., for the RGB format, the average of the mean values for red, green and blue). Since each infants of a group saw different exemplars of a category, for each infant, we created: an RDM representing signed size-differences for each pairwise comparison [(Size_right-image_−Size_left-image_)/(Size_right-image_+Size_left-image_)]; an RDM representing signed elongation-differences [(Elongation_right-image_−Elongation_left-image_)/(Elongation_right-image_+Elongation_left-image_)]; and an RDM representing color-differences in the form of Euclidean distance between the average color-value vector of two images. For each infant, we computed *Spearman* correlations between the DLT-RDM and each of the three RDMs. For each age group, individual correlation coefficients *ρ* were fisher-transformed and entered in a one-sample *t*-test (chance-level 0). For the signed RDMs (size and elongation), positive correlation values indicated longer looking times towards larger/more elongated objects.

#### Effects of size and elongation on categorization

As size and elongation correlated with the 4-month-olds’ DLTs in Experiment 1, and elongation correlated with 4-month-olds’ DLT in Experiments 1-2, we reassessed the effects of categorization after removing the variance explained by those physical properties of the images (Supplementary results 4). We performed a stepwise linear regression on the signed DLT-RDM of 4-month-olds, with size RDM and elongation RDM as regressors. Next, we performed the stepwise linear regression analysis on the absolute values of the residual matrices R [R = |signed DLT-RDM–β_size_ size-RDM – β_elongation_ elongation-RDM|].

#### Analysis of mean looking times (MLT)

For each group, we computed the MLT towards each category. Differences across categories were analyzed with a one-way repeated-measures ANOVA and followed up with pairwise *t*-tests.

### fMRI study on adults

#### Participants

Fifteen participants took part in the fMRI study (eight females; mean age: 24.9 years ± 3.6 *SD*). All had normal or corrected-to-normal vision, were screened for contraindications to fMRI, gave informed consent before participation and were paid for their time. The local ethics committee approved the study (CPP sud-est V).

#### Stimuli and experimental design

The fMRI study involved a main experiment on the same 72 object-stimuli (and eight categories) and a functional localizer session (see below). In the main experiment, all images were presented over two runs of 7.63 min each. Each run began with a warm-up block (10 s of fixation) and ended with a cool-down block (16 s of fixation), and included six sequences of eight blocks (one per category), for a total of 48 blocks of 6 s each (12 blocks per category in the whole experiment). Each block featured all nine exemplars of a category, in random order. Within a run, the inter-block interval duration was jittered (range: 0-6 s; total inter-block time per run: 144 s) to remove the overlap between estimates of the hemodynamic response. Jittering was optimized using the optseq tool of Freesurfer^64^. During a block, a blue cross was always present in the center of the screen, while stimuli appeared for 667 ms without interval. Participants were instructed to fixate the cross throughout the experiment, detect and report a change in the color of the cross (from blue to red in 40% of the blocks) by pressing a button with the right-index finger. This task was used to minimize eye movements and maintain vigilance. The main experiment lasted 15.26 min. Participants, lying down in the scanner, viewed the stimuli binocularly (~7° of visual angle) through a mirror above their head. Stimuli were back-projected onto a screen by a liquid crystal projector (frame rate: 60 Hz; screen resolution: 1024×768 pixels, screen size: 40×30 cm). For all the stimuli, the center of the image overlapped with the center of the screen. Stimulus presentation, response collection and synchronization with the scanner were controlled with the Psychtoolbox^65^ through MATLAB (MathWorks).

#### Functional localizer session

Stimuli and task were adapted from the fLoc package^66^. In this task, participants saw 180 grayscale photographs of five classes: body-stimuli (headless bodies and body parts), faces, places (houses and corridors), inanimate objects (cars and musical instruments) and scrambled objects. Stimuli were presented over two runs (5.27 min each). Runs began with a warm-up block (12 s) and ended with a cool-down block (16s), and included 72 blocks of 4 s each: 12 blocks for each object class with eight images per block (500 ms per image without interruption), randomly interleaved with 12 baseline blocks featuring an empty screen. To minimize low-level differences across categories, the view, size, and retinal position of the images varied across trials, and each item was overlaid on a 10.5° phase-scrambled background generated from another image of the set. During blocks of images, some images were repeated twice, interleaved by a different image. Participants pressed a button when they detected the repetition. Stimulus presentation, response collection and synchronization with the scanner were as in the main experiment.

#### Data acquisition

Imaging was conducted on a MAGNETOM-Prisma3T scanner (Siemens Healthcare). T2*-weighted functional volumes were acquired using a gradient-echo echo-planar imaging sequence (GRE-EPI; TR/TE: 2000/30 ms, flip angle: 80°, acquisition matrix: 96×92, FOV: 210×201, 56 transverse slices, slice thickness: 2.2 mm, no gap, multiband acceleration factor: 2 and phase encoding set to anterior/posterior direction). For the main experiment and the functional localizer session, we acquired four runs for a total of 790 frames per participant. Acquisition of high-resolution T1-weighted anatomical images was performed before the functional runs and lasted eight min (MPRAGE; TR/TE/TI: 3000/3.7/1100 ms, flip angle: 8°, acquisition matrix: 320×280, FOV: 256×224 mm, slice thickness: 0.8 mm, 224 sagittal slices, GRAPPA accelerator factor: 2). Acquisition of two field maps was performed at the beginning of the fMRI session.

#### Preprocessing

Functional images were preprocessed and analyzed using SPM12^67^ and the CoSMoMVPA toolbox (www.cosmomvpa.org) in combination with MATLAB. The first four volumes of each run were discarded, taking into account initial scanner gradient stabilization. Preprocessing of the remaining volumes involved despiking, slice time correction, geometric distortions correction using field maps, spatial realignment and motion correction using the first volume of each run as reference. Anatomical volumes were co-registered to the mean functional image, segmented into gray matter, white matter and cerebrospinal fluid in native space, and aligned to the probability maps in the Montreal Neurological Institute (MNI-152) as included in SPM12. The DARTEL method^68^ was used to create a flow field for each subject and an inter-subject template, which was registered in the MNI-152 space and used for normalization of functional images. Final steps included spatial smoothing (Gaussian kernel of 2 mm FWHM for main experiment and 6 mm FWHM for functional localizer), and removing of low-frequency drifts with a temporal high-pass filter (cutoff 128 s).

#### Analyses –Voxel Selection

The blood-oxygen-level-dependent (BOLD) signal of each voxel in each participant in the functional localizer task was modeled using five regressors, one for each of the five conditions, one regressor for baseline blocks, and six regressors of no interest for movement correction parameters. Within a gray matter mask based on the statistical map of the second-level (group) analysis, voxels that responded to visual object stimulation across the whole brain were selected with the contrast ([bodies+faces+places+objects+scrambled objects] > [baseline]) (voxelwise *P* = 0.05).

#### ROIs

We created a model of visual object categorization based on the neural activity patterns evoked by the eight visual object categories, in three ROIs across the visual cortex. Voxels selected with the functional localizer task were divided in three sectors, using three bilateral masks of the early visual cortex (EVC), ventral occipitotemporal cortex (VOTC) and lateral occipitotemporal cortex (LOTC), extracted from the SPM Anatomy toolbox^69^. The EVC included all voxels that responded to visual object stimulation during the functional localizer task and fell within early visual areas V1/V2 or ventral/dorsal extrastriate areas V3v/V3d. The VOTC-ROI was defined within a mask encompassing the extrastriate area V4v and fusiform gyrus (FG1/FG2/FG3/FG4). The LOTC-ROI included voxels within a mask encompassing lateral occipital areas V4la/V4lp and V5.

#### Vector-of-ROIs

To capture more local effects of visual categorization, we created RDMs from neural patterns evoked by the eight objects categories, at each consecutive partition along the antero-posterior axis that forms the visual ventral stream. First, a mask of the visual ventral stream was defined by selecting all the voxels that responded to visual object presentation in the functional localizer task and fell within V1/V2, V3v/V4v and FG1/FG2/FG3/FG4. This mask was then divided into 38 consecutive slices of 2.2 mm width, along the antero-posterior axis.

#### fMRI-based RDMs

Independently for each run of the main fMRI experiment, the BOLD signal of each voxel in each participant was modeled in a general linear model, using eight regressors for the eight object-categories, one regressor for baseline fixation blocks, and six regressors of no interest for movement correction parameters. For each voxel in the brain, groupwise beta values for a given category were extracted in a second-level group analysis using the contrast [category > baseline]. Finally, for each ROI and for each slice along the vector of ROIs, we computed an RDM representing dissimilarities between neural patterns (one minus *Pearson* coefficient *r*) for each between-category and within-category comparison (Fig. 3b).

#### RSA

For each RDM extracted from the three ROIs (EVC, VOTC, and LOTC), we computed the correlation with the DLT-RDM of each individual infant (*Spearman* correlation). Correlation coefficients *ρ* of all infants of an age group were fisher-transformed and entered in a one-sample *t* test (two-tailed), to assess the relationship between the infants’ looking behavior and object representations in different sectors of the adults’ visual cortex. The above analysis was repeated using RDMs derived from each slice of the vector of ROIs. Clusters of consecutive slices showing correlation with the infants’ DLT-RDMs were identified as follows: for each slice, the individual’s fisher-transformed coefficients *ρ* were entered into a one-sample *t* test; the *t*-values for each slice were transformed in *z*-values and analyzed in a cluster mass test^70^ with a threshold of *z* = 1.64 (*P* = 0.05, two-tailed; Fig. 3d) and 1000 permutations.

## ACKNOWLEDGEMENTS

Funding for this project was provided by a European Research Council Starting Grant awarded to L.P. (Project: THEMPO, Grant Agreement 758473), by a Fyssen Foundation research Grant awarded to J.R.H. and by an Agence Nationale pour la Recherche (ANR) Grant awarded to J.R.H. (ANR-16-CE28-0006-TACTIC).

We would like to acknowledge Marion Gasselin for infants’ recruitment.

## AUTHOR CONTRIBUTIONS

C.S., J.R.H. and L.P. designed research; C.S. and E.A. performed research; all authors contributed analytic tools and data analysis; C.S., J.R.H. and L.P. wrote the paper.

## DATA AVAILABILITY

Group fMRI and eye-tracking are available at the Open Science Framework repository for this project (https://osf.io/6rm7a/?view_only=dcd418e45e074e379edee09ba36840be).

Fig. 2: raw data available at https://osf.io/6rm7a/?view_only=dcd418e45e074e379edee09ba36840be

Fig. 3: raw data available at …

## CODE AVAILABILITY

Code for the main analyses is available at the Open Science Framework repository for this project (https://osf.io/6rm7a/?view_only=dcd418e45e074e379edee09ba36840be).

