## Supplementary for "Visual object categorization in infancy"

### Supplementary Results

#### 1. Stepwise linear regression analysis (Experiments 1-2)

We run a stepwise linear regression with the six synthetic models as regressors for each group in Experiments 1-2. As reported in the main text, this analysis showed an effect of the eight-category model, animacy model, and humanness model for 19-month-old infants (Experiment 1), and of the animacy model only for 10-month-old infants (Experiment 1) and 4-month-old infants in Experiment 2. See Supplementary Table 1 for statistical values of this analysis.

**Supplementary Table 1. Results of representational similarity analysis (Stepwise linear regression) reflecting relationships between the infants' DLT-RDMs and the synthetic models of categorization**

| Exp. | Age | Regressor | Mean $\beta$ *(SD)* | CI (min – max) | *t* (df) | *P* | Cohen's *d* |
|---|---|---|---|---|---|---|---|
| 1 | 19 m | **Eight-categories** | **.090 *(.112)*** | **.026 – .155** | **4.023 (24)** | **<.001** | **.804** |
| | | **Animacy** | **.077 *(.109)*** | **.014 – .139** | **3.514 (24)** | **.002** | **.702** |
| | | **Humanness** | **.133 *(.215)*** | **.009 – .256** | **3.091 (24)** | **.005** | **.618** |
| | | Faces/Bodies | .022 *(.135)* | -.056 – .099 | 0.811 (24) | *n.s.* | .163 |
| | | Natural/Artificial | .017 *(.219)* | -.109 – .143 | 0.384 (24) | *n.s.* | .078 |
| | | Big/Small | .066 *(.176)* | -.036 – .167 | 1.866 (24) | .074 | .375 |
| 1 | 10 m | Eight-categories | .051 *(.113)* | -.015 – .118 | 2.222 (23) | .036 | .451 |
| | | **Animacy** | **.059 *(.098)*** | **.002 – .117** | **2.981 (23)** | **.007** | **.608** |
| | | Humanness | .002 *(.200)* | -.116 – .119 | 0.039 (23) | *n.s.* | .010 |
| | | Faces/Bodies | .045 *(.181)* | -.062 – .152 | 1.207 (23) | .240 | .249 |
| | | Natural/Artificial | -.026 *(.122)* | -.098 – .046 | -1.054 (23) | *n.s.* | .213 |
| | | Big/Small | .038 *(.128)* | -.038 – .113 | 1.448 (23) | .161 | .297 |
| 1 | 4 m | Eight-categories | .013 *(.101)* | -.047 – .073 | 0.634 (23) | *n.s.* | .129 |
| | | Animacy | .014 *(.118)* | -.055 – .084 | 0.598 (23) | *n.s.* | .119 |
| | | Humanness | .034 *(.206)* | -.088 – .156 | 0.809 (23) | *n.s.* | .165 |
| | | Faces/Bodies | .012 *(.200)* | -.106 – .130 | 0.304 (23) | *n.s.* | .060 |
| | | Natural/Artificial | .031 *(.250)* | -.117 – .178 | 0.606 (23) | *n.s.* | .124 |
| | | Big/Small | .021 *(.246)* | -.124 – .166 | 0.421 (23) | *n.s.* | .085 |
| 2 | 4 m | Eight-categories | .026 *(.142)* | -.057 – .110 | 0.906 (23) | *n.s.* | .183 |
| | | **Animacy** | **.074 *(.098)*** | **.016 – .132** | **3.697 (23)** | **.001** | **.755** |
| | | Humanness | -.061 *(.223)* | -.192 – .070 | -1.341 (23) | .193 | .274 |
| | | Faces/Bodies | .019 *(.211)* | -.106 – .143 | 0.430 (23) | *n.s.* | .090 |
| | | Natural/Artificial | .096 *(.296)* | -.079 – .271 | 1.591 (23) | .125 | .324 |
| | | Big/Small | .008 *(.232)* | -.129 – .145 | 0.165 (23) | *n.s.* | .034 |

*Note: Exp., experiment; m, months; CI, 99.17% confidence interval; Highlighted in bold are the significant results; $\alpha$ = .0083, two-tailed; n.s.= non-significant results with P > .250.*

**2. Difference between within-category and between-category DLTs (Experiments 1-2).**

For each Experiment, for each age group, for each of the six categorization models, we tested whether within-category differential looking times (DLTs) were lower than between-category DLTs. As mentioned in the main text, this analysis showed an effect of the eight-category model, animacy model, and humanness model in 19-month-old infants (Experiment 1), and of the animacy model only in 10-month-old infants (Experiment 1) and in 4-month-old infants of Experiment 2. Supplementary Table 2 report all statistical values of this analysis.

**Supplementary Table 2. Results of the DLTs analyses of within *versus* between category comparisons**

| Exp. | Age | Comparisons within/between | Mean of the difference *(SD)* | CI (min – max) | *t* (df) | *P* | Cohen's *d* |
|---|---|---|---|---|---|---|---|
| 1 | 19 m | **Diagonal** | **-.135 *(.085)*** | **-inf – -.091** | **-7.967 (24)** | **<.0001** | **1.588** |
| | | **Animate and inanimate** | **-.095 *(.106)*** | **- inf – -.041** | **-4.481 (24)** | **<.0001** | **.896** |
| | | **Human and nonhuman** | **-.158 *(.190)*** | **- inf – -.060** | **-4.160 (24)** | **<.001** | **.832** |
| | | Faces and Bodies | -.057 *(.141)* | - inf – .016 | -2.013 (24) | .028 | .404 |
| | | Natural and Artificial | -.035 *(.210)* | - inf – .074 | -.825 (24) | .209 | .167 |
| | | Big and Small | -.074 *(.178)* | - inf – .018 | -2.071 (24) | .025 | .416 |
| 1 | 10 m | Diagonal | -.057 *(.137)* | - inf – .016 | -2.018 (23) | .028 | .416 |
| | | **Animate and inanimate** | **-.061 *(.103)*** | **- inf – -.007** | **-2.919 (23)** | **.004** | **.592** |
| | | Human and nonhuman | -.003 *(.202)* | - inf – .104 | -.066 (23) | *n.s.* | .015 |
| | | Faces and Bodies | -.045 *(.202)* | - inf – .061 | -1.096 (23) | .142 | .223 |
| | | Natural and Artificial | .029 *(.139)* | -inf – .102 | 1.036 (23) | *n.s.* | .209 |
| | | Big and Small | -.041 *(.145)* | - inf – .036 | -1.370 (23) | .092 | .283 |
| 1 | 4 m | Diagonal | -.048 *(.120)* | - inf – .016 | -1.940 (23) | .032 | .400 |
| | | Animate and inanimate | -.022 *(.121)* | - inf – .042 | -.873 (23) | .196 | .182 |
| | | Human and nonhuman | -.036 *(.216)* | -inf – .078 | -.811 (23) | .213 | .167 |
| | | Faces and Bodies | -.013 *(.205)* | - inf – .095 | -.316 (23) | *n.s.* | .063 |
| | | Natural and Artificial | -.043 *(.252)* | - inf – .090 | -.840 (23) | .205 | .171 |
| | | Big and Small | -.027 *(.262)* | - inf – .111 | -.513 (23) | *n.s.* | .103 |
| 2 | 4 m | Diagonal | -.024 *(.133)* | - inf – .046 | -.891 (23) | .191 | .180 |
| | | **Animate and inanimate** | **-.076 *(.104)*** | **- inf – -.021** | **-3.583 (23)** | **<.001** | **.731** |
| | | Human and nonhuman | .087 *(.232)* | - inf – .209 | 1.836 (23) | *n.s.* | .375 |
| | | Faces and Bodies | .018 *(.225)* | - inf – .136 | .383 (23) | *n.s.* | .080 |
| | | Natural and Artificial | -.086 *(.308)* | - inf – .077 | -1.360 (23) | .094 | .279 |
| | | Big and Small | -.014 *(.259)* | - inf – .123 | -.267 (23) | *n.s.* | .054 |

*Note: Exp., experiment; m, months; CI, 99.17% confidence interval; Highlighted in bold are the significant results; α = .0083, one-tailed; n.s.= non-significant results with P > .250.*

#### 3. Analysis of mean looking times (Experiments 1-2)

Separate one-way ANOVAs for 4-, 10- and 19-month-olds in Experiment 1 and for 4-month-olds in Experiment 2, revealed the effect of categories on mean looking times (MLTs). The following post-hoc analyses were carried out to follow up on the effect of category. For 19-month-old infants in Experiment1, pairwise comparisons (one sample *t*-tests, $\alpha_{corrected}$: 0.0018) revealed longer looking times towards animate than inanimate objects (9 out of 16 comparisons were significant; 4 of the 7 remaining comparisons showed non-significant trends, that is, effects that did not reach the significance level, corrected for multiple comparisons). A *t*-test comparing all animate *vs.* all inanimate categories confirmed this effect (animate: $M \pm SD$ = 2.103 s ± 0.344; inanimate: $M \pm SD$ = 1.598 s ± 0.247; $M_{difference} \pm SD$ = 0.505 ± 0.335; 95% CI = 0.367 – 0.643; $t(24) = 7.551$, $P < 0.0001$; $d = 1.509$). Comparisons between human and nonhuman categories showed significant or non-significant trends ($Ps < .03$), which can be summarized as a preference for nonhuman categories ($M \pm SD$ = 2.317 s ± 0.463) over human ($M \pm SD$ = 1.889 s ± 0.452; $M_{difference} \pm SD$ = -0.428 ± 0.602; 95% CI = -0.676 – -0.179; $t(24) = -3.552$, $P = 0.002$; $d = 0.711$). Among the inanimate categories, only the comparison between natural-small *vs.* artificial-big showed a non-significant trend ($P = 0.019$; for other comparisons all $Ps > 0.11$).

The same analyses on 10-month-old infants (Experiment 1) showed that infants looked longer at the animate than inanimate categories (10 out of 16 comparisons were significant with $\alpha_{corrected} = 0.0018$; 4 of the 6 remaining comparisons showed non-significant trends with $Ps < 0.05$; for the remaining two comparisons, $Ps > 0.062$). A *t*-test comparing MLTs for all animate *vs.* all inanimate objects showed significantly longer MLTs for animate ($M \pm SD$ = 2.044 s ± 0.316) than for inanimate categories ($M \pm SD$ = 1.601 s ± 0.263; $M_{difference} \pm SD$ = 0.443 ± 0.314; 95% CI = 0.310 – 0.576; $t(23) = 6.901$, $P < 0.0001$; $d = 1.409$). When two animate or two inanimate categories were compared, only 1 out of 12 comparisons showed a non-significant trend ($P = 0.007$; for all other comparisons $Ps > 0.056$).

For 4-month-old infants in Experiment 1, pairwise comparisons (one sample *t*-tests, $\alpha_{corrected}$ = 0.0018, two-tailed) revealed that infants looked longer at human faces relative to all other categories (*Ps* < 0.001 for 6 of the 7 comparisons between the human face category and other categories respectively), except the natural big objects, for which the MLTs only marginally differed (*P* = 0.002). No other difference was observed within the animate categories. Within the inanimate categories, infants looked longer at big than small objects, whether artificial or natural (*Ps* < 0.0001).

Four-month-old infants in Experiment 2 showed to prefer (i.e., looked longer at) human faces *vs.* all other categories (all *Ps* < 0.0001) and big *vs.* small (natural or artificial) inanimate objects (all *Ps* < 0.001). Unlike in Experiment 1, the preference for human faces and big objects in Experiment 2 was not conflated with the preference for large images, as all images featured the same number of pixels. However, the mean looking time for individual images, averaged across participants, negatively correlated with the image shape elongation ($\rho = -0.396$, $p < 0.001$): the less elongated the shape, the longer the looking time (see also Supplementary Results 4). See Supplementary Table 3 for *t*-values and *P*-values of all the pairwise comparisons reported here.

**4. Relationship between MLTs and low-level features of images.**

Because 4-month-olds' looking behavior appeared to be guided by low-level features such as the size of images or the elongation (Supplementary Fig. 1), we tested whether the categories of images that infants looked at for a longer time, corresponded to the categories including larger images and/or less elongated objects. To this end, we computed the MLTs, the mean number of pixels and the mean elongation ratio for each category (Supplementary Fig. 2). The three preferred categories, human faces and natural/artificial big objects, constituted the largest (Experiment 1) and least elongated (Experiments 1-2) images. Furthermore, mean looking times (MLTs) correlated significantly with both size and elongation in Experiment 1, and with elongation in Experiment 2 (see Main text).

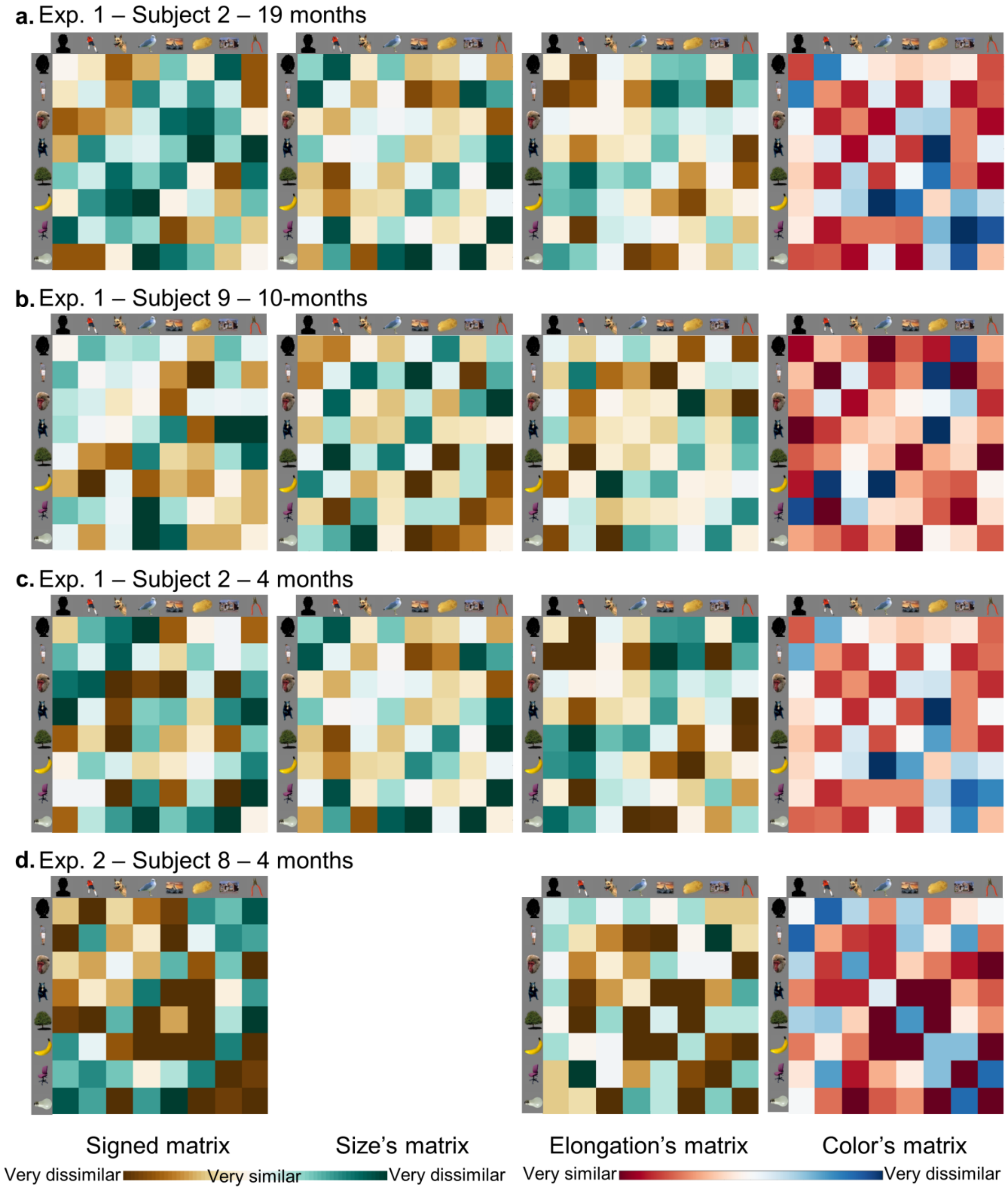


**Supplementary Figure 1. Signed DLT-RDM (left), size RDM (middle left), elongation RDM (middle right) and color RDM (right) for 4-, 10- and 19-month-olds in Experiment 1 and 4-month-olds in Experiment 2**. Example of a signed DLT-RDM, a size RDM, an elongation RDM and a color RDM for one subject of the 19-month-old group (**a**), the 10-month-old group (**b**) and the 4-month-old group (**c**) in Experiment 1, and one subject of the 4-month-old group in Experiment 2 (**d**). Only silhouette of the face-stimuli used in the experiment are shown, because verification of consent of people in the pictures is incompatible with the rapid and automated nature of preprint posting.

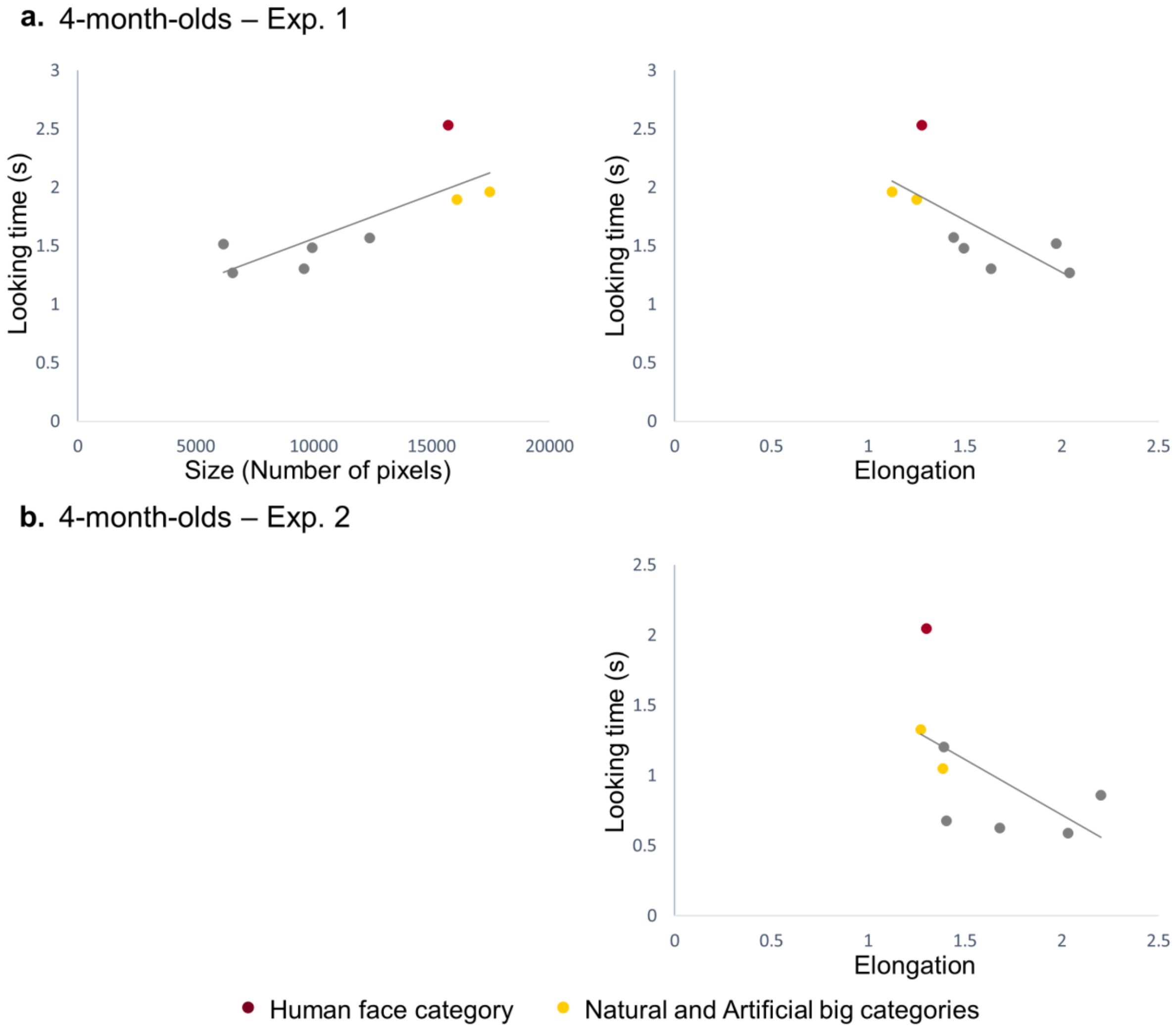


**Supplementary Figure 2. Looking times as a function of image size (left) and elongation (right) for 4-month-olds in Experiments 1-2.** Relationship between looking times (s) and average size of the images (number of pixels) for each category and average elongation (ratio between width and height) for each category for 4-month-olds in Experiment 1 (**a**) and relationship between looking times (s) and average elongation for each category for 4-month-olds in Experiment 2 (**b**).

Given these results, for each group of 4-month-olds, we performed a novel stepwise linear regression analysis and DLTs analysis, after removing the variance explained by size and elongation in Experiment 1, and by elongation only in Experiment 2 (Supplementary Tables 4- 5).

**Supplementary Table 4. Results of the stepwise linear regression with artificial models, removing variance explained by size and/or elongation of 4-month-old infants' DLT-RDM in experiment 1 and 2**

| Exp. | Regressor | Mean *β* *(SD)* | CI (min – max) | *t* (df) | *P* | Cohen's *d* |
|---|---|---|---|---|---|---|
| 1 | Eight-categories | -.002 *(.101)* | -.062 – .057 | -.117 (23) | *n.s.* | .020 |
| | Animacy | .012 *(.142)* | -.072 – .096 | .416 (23) | *n.s.* | .085 |
| | Humanness | .045 *(.246)* | -.100 – .189 | .887 (23) | *n.s.* | .183 |
| | Faces/Bodies | -.026 *(.237)* | -.166 – .114 | .531 (23) | *n.s.* | .110 |
| | Natural/Artificial | -.029 *(.186)* | -.139 – .081 | .755 (23) | *n.s.* | .156 |
| | Big/Small | -.042 *(.179)* | -.148 – .064 | 1.148 (23) | *n.s.* | .235 |
| 2 | Eight-categories | -.020 *(.121)* | -.091 – .051 | -.806 (23) | *n.s.* | .165 |
| | **Animacy** | **.064 *(.096)*** | **.008 – .121** | **3.282 (23)** | **.003** | **.667** |
| | Humanness | -.069 *(.212)* | -.194 – .056 | -1.584 (23) | .127 | .325 |
| | Faces/Bodies | -.013 *(.204)* | -.133 – .108 | -.303 (23) | *n.s.* | .064 |
| | Natural/Artificial | .085 *(.285)* | -.084 – .253 | 1.453 (23) | .160 | .298 |
| | Big/Small | -.016 *(.215)* | -.142 – .111 | -.365 (23) | *n.s.* | .074 |

*Note: Exp., experiment; CI, confidence interval; Highlighted in bold are the significant results; α = .0083, two-tailed; n.s.= non-significant results with P > .250.*

**Supplementary Table 5. Results of the DLTs analysis of within *versus* between category comparisons, removing variance explained by size and/or elongation of 4-month-old infants in experiment 1 and 2**

| Exp. | Comparisons within/between | Mean of the difference *(SD)* | CI (min – max) | *t* (df) | *P* | Cohen's *d* |
|---|---|---|---|---|---|---|
| 1 | Diagonal | -.021 *(.110)* | - inf – .037 | -.943 (23) | .178 | .191 |
| | Animate and inanimate | -.016 *(.143)* | - inf – .059 | -.543 (23) | *n.s.* | .112 |
| | Human and nonhuman | -.048 *(.247)* | - inf – .082 | -.955 (23) | .175 | .194 |
| | Faces and Bodies | .015 *(.238)* | - inf – .141 | .310 (23) | *n.s.* | .063 |
| | Natural and Artificial | .041 *(.203)* | - inf – .148 | .998 (23) | *n.s.* | .202 |
| | Big and Small | .046 *(.179)* | - inf – .141 | 1.257 (23) | *n.s.* | .257 |
| 2 | Diagonal | .002 *(.119)* | - inf – .064 | .075 (23) | *n.s.* | .017 |
| | **Animate and inanimate** | **-.068 *(.101)*** | **- inf – -.015** | **-3.314 (23)** | **.002** | **.673** |
| | Human and nonhuman | .090 *(.226)* | - inf – .209 | 1.937 (23) | *n.s.* | .398 |
| | Faces and Bodies | .043 *(.221)* | - inf –.160 | .959 (23) | *n.s.* | .195 |
| | Natural and Artificial | -.079 *(.307)* | - inf – .083 | -1.256 (23) | .111 | .257 |
| | Big and Small | .018 *(.248)* | - inf – .149 | .355 (23) | *n.s.* | .073 |

*Note: Exp., experiment; CI, confidence interval; Highlighted in bold are the significant results; α = .0083, two-tailed; n.s. = non-significant results with P > .250*

**5. Between-group comparisons**

All analyses reported in the main text and in the supplementary material so far concur to identify developmental changes in infants' visual categorization. In Experiment 1, categorization by animacy was observed in 10- and 19-month-olds, but not in 4-month-olds. Categorization by humanness and additional categories represented in the eight-category model was observed in 19-month-olds but not in 4- and 10-month-olds.

To further test these developmental differences, for each of the above categorization models that appeared different across groups (animacy, humanness and eight-category model), we analyzed the variation of the mean difference between averaged between-categories *vs.* averaged within-category DLTs with a one-way ANOVA, with Age as between-subject factor (4 months, 10 months, 19 months).

As for the animacy model, we found a trend for the effect of Age ($F(2,70) = 2.752$; $P = 0.071$; $\eta^2 = 0.073$). Pairwise comparisons showed that 4-month-olds differed significantly from 19-month-olds (4-month-olds: $M \pm SD = -0.022 \pm 0.121$; 19-month-olds: $M \pm SD = -0.095 \pm 0.106$; 95% CI = 0.009 – 0.139; $t(47) = 2.273$; $P = 0.028$; $d = 0.649$) but not from 10-month-olds (10-month-olds: $M \pm SD = -0.061 \pm 0.103$; 95% CI = -0.026 – 0.105; $t(46) = 1.225$; $P = 0.227$; $d = 0.354$). Ten- and 19-month-olds did not differ (95% CI = -0.026 – 0.094; $t(47) = 1.144$; $P = 0.258$; $d = 0.328$).

As for the humanness model, the effect of Age was significant ($F(2,70) = 4.023$; $P = 0.022$; $\eta^2 = 0.103$). Nineteen-month-olds differed from both 4-month-olds (4-month-olds: $M \pm SD = -0.036 \pm 0.216$; 19-month-olds: $M \pm SD = -0.158 \pm 0.190$; 95% CI = 0.006 – 0.239; $t(47) = 2.109$; $P = 0.040$; $d = 0.602$) and 10-month-olds (10-month-olds: $M \pm SD = -0.003 \pm 0.202$; 95% CI = 0.043 – 0.268; $t(47) = 2.780$; $P = 0.008$; $d = 0.794$). Four- and 10-month-olds did not differ (95% CI = -0.155 – 0.088; $t(46) = -0.548$ ; $P = 0.586$; $d = 0.158$).

As for the eight-categories model the effect of Age was significant ($F(2,70) = 4.292$; $P = 0.018$; $\eta^2 = 0.109$). Nineteen-month-olds differed from both 4- (4-month-olds: $M \pm SD = -0.048 \pm$

0.120; 19-month-olds: $M \pm SD$ = -0.135 ± 0.085; 95% CI = 0.028 – 0.147; $t(47) = 2.966$; $P = 0.005$; $d = 0.844$) and 10-month-olds (10-month-olds: $M \pm SD$ = -0.057 ± 0.137; 95% CI = 0.013 – 0.144; $t(47) = 2.426$; $P = 0.019$; $d = 0.690$). Four- and 10-month-olds did not differ (95% CI = -0.066 – 0.084; $t(46) = 0.243$; $P = 0.809$; $d = 0.070$).

In sum, categorization by humanness and by the so-called eight-categories model changed and developed between 10 and 19 months. Categorization by animacy differed between 4 and 19 months, but not between 4 and 10 months.

**6. Experiment 2: Data analysis with more lenient inclusion criteria**

In Experiment 2, the attrition rate for 4-month-olds was larger compared to 4-month-olds in Experiment 1 (37% in Exp. 1 and 50% in Exp. 2). Data from 24 out of 48 tested infants were discarded in Experiment 2, while data from 14 out of 38 tested infants were discarded in Experiment 1. A larger attrition rate may lead to the inclusion of more attentive infants, which could explain the different results in Experiment 2 compared to Experiment 1.

To take into account this possibility, we ran a novel analysis on the data from Experiment 2, using inclusion criteria that yielded an attrition rate comparable to that of Experiment 1. In particular, we modified the criterion for trial inclusion, selecting all trials in which infants looked at images for at least 800 ms (insteasd of 1000 ms in the original analysis reported in the main text). This change yielded the exclusion of 16 out of 48 infants (attrition rate of 33%). The analysis of this dataset confirmed the results reported in the main text. The stepwise linear regression analyzing the structure of the absolute RDMs identified only one significant regressor ($\alpha_{corrected} = 0.0083$), corresponding to the animacy model (mean $\beta \pm SD$ = 0.066 ± 0.094; 99.17% CI = 0.020 – 0.113; $t(31) = 3.991$; $P < 0.001$; $d = 0.705$) (for other regressors, $ts < 2.313$; $Ps > 0.028$). The DLTs analysis also showed that the difference between within-category and between-category DLTs was only significant for the animacy model ($M_{difference} \pm SD$ = -0.065 ± 0.094; 99.17% CI = -inf – -0.023; $t(31) = -3.927$; $P < 0.001$; $d = 0.691$) (for other models, $ts > -2.208$; $Ps > 0.017$).

**Supplementary Table 3. Absolute t-values and P-values for pairwise comparisons (t tests) of MLTs**

| Age | Categories | HB | | NHF | | NHB | | NB | | NS | | AB | | AS | |
|---|---|---|---|---|---|---|---|---|---|---|---|---|---|---|---|
| | | *t* | *P* | *t* | *P* | *t* | *P* | *t* | *P* | *t* | *P* | *t* | *P* | *t* | *P* |
| 19 m | HF | 1.083 | *n.s.* | **4.083** | **<0.001** | 3.079 | 0.005 | 2.445 | 0.022 | 3.003 | 0.006 | 1.187 | 0.247 | 1.921 | 0.067 |
| | HB | | | 3.138 | 0.005 | 2.317 | 0.029 | **3.706** | **0.001** | 3.461 | 0.002 | 1.903 | 0.069 | 2.567 | 0.017 |
| | NHF | | | | | 0.472 | *n.s.* | **6.875** | **<0.0001** | **8.575** | **<0.0001** | **6.005** | **<0.0001** | **5.914** | **<0.0001** |
| | NHB | | | | | | | **5.624** | **<0.0001** | **8.298** | **<0.0001** | **4.702** | **<0.0001** | **5.553** | **<0.0001** |
| | NB | | | | | | | | | 1.264 | 0.219 | 1.076 | *n.s.* | 0.271 | *n.s.* |
| | NS | | | | | | | | | | | 2.505 | 0.019 | 1.650 | 0.112 |
| | AB | | | | | | | | | | | | | 0.800 | *n.s.* |
| 10 m | HF | 1.152 | *n.s.* | 0.564 | *n.s.* | 1.830 | 0.080 | **5.528** | **<0.0001** | **4.148** | **<0.001** | **3.779** | **<0.001** | **6.163** | **<0.0001** |
| | HB | | | 0.452 | *n.s.* | 1.050 | *n.s.* | **3.717** | **0.001** | 2.974 | 0.007 | 2.517 | 0.019 | **5.050** | **<0.0001** |
| | NHF | | | | | 1.566 | 0.131 | **3.642** | **0.001** | **3.701** | **0.001** | 2.878 | 0.009 | **4.918** | **<0.0001** |
| | NHB | | | | | | | 2.737 | 0.012 | 1.959 | 0.062 | 1.404 | 0.174 | **4.287** | **<0.001** |
| | NB | | | | | | | | | 0.221 | *n.s.* | 1.627 | 0.117 | 2.012 | 0.056 |
| | NS | | | | | | | | | | | 0.786 | *n.s.* | 1.848 | 0.078 |
| | AB | | | | | | | | | | | | | 2.972 | 0.007 |
| 4 m | HF | **5.878** | **<0.0001** | **6.702** | **<0.0001** | **6.887** | **<0.0001** | 3.517 | 0.002 | **8.056** | **<0.0001** | **4.115** | **<0.001** | **7.954** | **<0.0001** |
| | HB | | | 0.457 | *n.s.* | 0.497 | *n.s.* | **5.563** | **<0.0001** | 1.872 | 0.074 | 3.033 | 0.006 | 2.945 | 0.007 |
| | NHF | | | | | 0.716 | *n.s.* | 3.404 | 0.002 | 1.836 | 0.079 | 2.321 | 0.030 | 2.855 | 0.009 |
| | NHB | | | | | | | **5.470** | **<0.0001** | 1.931 | 0.066 | 3.409 | 0.002 | 2.769 | 0.011 |
| | NB | | | | | | | | | **5.934** | **<0.0001** | 0.546 | *n.s.* | **6.944** | **<0.0001** |
| | NS | | | | | | | | | | | **4.552** | **<0.001** | 0.357 | *n.s.* |
| | AB | | | | | | | | | | | | | **4.416** | **<0.001** |
| 4 m | HF | **10.673** | **<0.0001** | **8.143** | **<0.0001** | **13.239** | **<0.0001** | **8.214** | **<0.0001** | **14.461** | **<0.0001** | **7.712** | **<0.0001** | **13.536** | **<0.0001** |
| | HB | | | **4.011** | **<0.001** | 2.979 | 0.007 | 1.745 | 0.094 | **3.837** | **<0.001** | **4.684** | **<0.001** | **3.752** | **0.001** |
| | NHF | | | | | **7.995** | **<0.0001** | 1.995 | 0.058 | **5.929** | **<0.0001** | 1.446 | 0.162 | **7.975** | **<0.0001** |
| | NHB | | | | | | | **4.070** | **<0.001** | 0.999 | *n.s.* | **7.173** | **<0.0001** | 1.297 | 0.207 |
| | NB | | | | | | | | | **4.305** | **<0.001** | 3.425 | 0.002 | **5.914** | **<0.0001** |
| | NS | | | | | | | | | | | **8.650** | **<0.0001** | 0.048 | *n.s.* |
| | AB | | | | | | | | | | | | | **9.041** | **<0.0001** |

*Note: Exp., experiment; m, months; HF: Human Face; HB: Human Body; NHF: NonHuman Face; NHB: NonHuman Body; NB: Natural Big; NS: Natural Small; AB: Artificial Big; AS: Artificial Small; Highlighted in bold are the significant results; α = .0018; n.s.= non-significant with P > .250*
